# Oxycodone self-administration and reinstatement in male and female squirrel monkeys: Effects of alternative reinforcer availability

**DOI:** 10.1101/2023.01.12.523850

**Authors:** Fernando B de Moura, Raymond G Booth, Stephen J Kohut

## Abstract

The use of non-drug alternative reinforcers has long been utilized as a component of therapeutic interventions for the management of substance use disorder; however, the conditions under which alternative reinforcers are most effective are not well characterized. This study evaluated the impact of varying the magnitude of an alternative reinforcer on oxycodone self-administration and reinstatement in male and female squirrel monkeys. Subjects (n=4/sex) were trained under concurrent second-order schedules of reinforcement for intravenous oxycodone (0.001-0.1mg/kg/inj) on one lever, and sweetened condensed milk (5, 10, 20, 30% in water) on another. Oxycodone-primed reinstatement was evaluated by administering 0.32mg/kg oxycodone prior to sessions in which saline was available on the drug-paired lever. During oxycodone self-administration sessions, milk availability decreased oxycodone self-administration and preference in a concentration-dependent manner; low milk concentrations were more effective at decreasing oxycodone’s reinforcing potency in males. During reinstatement tests, milk significantly attenuated oxycodone-primed responding in both males and females; low milk concentrations were more effective at decreasing the priming effects of oxycodone in females. That alternative reinforcers differentially impacted self-administration and reinstatement in a sex-dependent manner suggests that treatment strategies that utilize alternative reinforcers may be more effective in males or females depending on when they are implemented.

## Introduction

Substance use disorders, such as opioid use disorder (OUD), are influenced by a combination of pharmacological and behavioral mechanisms, and, as a result, are inherently complex to manage. Treatment methodologies for OUDs include pharmacological (e.g., methadone maintenance, Suboxone^®^) and behavioral (e.g., 12-step programs, contingency management, voucher programs); however, despite varying levels of success, relapse to drug use frequently occurs and increases in both the abuse and relapse of opioids have dramatically increased since the onset of the COVID-19 pandemic [1,2,3]. Therefore, identifying either novel treatment modalities and/or improving upon existing OUD management therapies remains a critical need.

The use of non-drug reinforcer alternatives is a key component of therapeutic interventions such as contingency management and voucher programs [4,5,6]. Such strategies have been relatively successful in humans, as cocaine, nicotine, alcohol, and heroin use has been shown to significantly decrease when money is available as an alternative [7,8,9,10,11]. Therefore, a more comprehensive understanding of how both drug and non-drug reinforcer magnitudes influence OUD-related behaviors may identify conditions necessary to increase the effectiveness of alternative reinforcers in managing OUD.

Intravenous drug self-administration procedures in which subjects have concurrent access to drug and non-drug reinforcers during experimental sessions – e.g., drug vs. food “choice” procedures - have been utilized to evaluate the impact of alternative reinforcers on the abuse-related effects of drugs and are thought to be translatable to the human condition [7,8,9,12,13,14]. In preclinical studies, a concurrently available palatable food reinforcer has been shown to decrease intake of various drugs in both rodents and nonhuman primates, including prescription opioids [15]. Although most studies use a single alternative reinforcer with a fixed magnitude, Nader and Woolverton [16] demonstrated that increasing the magnitude of an alternative reinforcer (e.g., number of food pellets) decreases the frequency in which cocaine is self-administered in nonhuman primates. That result was consistent with experiments in humans in which increasing the amount of a monetary reinforcer systematically decreased opioid choice [4,5]. However, another study found that the availability of an alternative reinforcer is less effective at attenuating fentanyl self-administration in male rats compared to female rats [15], suggesting that alternative reinforcers may be qualitatively and/or quantitatively different in magnitude and effectiveness between the sexes.

Although the magnitude of an alternative reinforcer has been shown to be an important variable for decreasing drug self-administration, the importance of reinforcer magnitude on relapse-related drug-seeking is largely unknown. Studies in rodents have shown that reinforcer density (i.e., the frequency of reinforcer presentations) can attenuate reinstatement of drug-seeking following extinction. For example, Craig et al. [17] demonstrated that during extinction training, rats exposed to a schedule of palatable food reinforcement were less likely to maintain responding on a lever previously paired with i.v. cocaine injections. Furthermore, rats that were food-trained during extinction with a low ratio requirement were less likely to perseverate on a drug-associated lever than those trained on a high ratio requirement [17]. These results support the hypothesis that alternative reinforcers can attenuate drug-seeking behaviors and that they can be systematically manipulated to increase their effectiveness in curtailing abuse-related behaviors.

The present study was designed to evalute the impact of changing the magnitude of a concurrently available alternative reinforcer on oxycodone self-administration and reinstatement in nonhuman primates. Investigating the influence of alternative reinforcers on both self-adminstration and reinstatement behaviors allows for the characterization of a dynamic range of abuse-related behaviors- from actively engaging in drug use (i.e., drug-taking) to relapse-related behaviors and searching for drug availability (i.e., drug-seeking). Further, a growing body of clinical, epidemiological, and laboratory-based evidence indicates distinct sex-dependent patterns of opioid misuse [18,19,20,21,22,23,24,25,26]. Thus, sex as a biological variable was considered to determine whether alternative reinforcers were similarly effective in modulating the abuse related effects of oxycodone in male and female subjects. Results from this study may provide insights for using alternative reinforcers as a component of treatment for the management of OUD.

## Methods

### Subjects

Eight adult squirrel monkeys (4 male/4 female; *saimiri sciureus*), weighing 750-1000g, were studied in daily experimental sessions (Mon-Fri). All subjects were pair-housed in stainless steel cages located in a climate-controlled vivarium under an automated 12-h light/dark cycle. Subjects had ad libitum access to water and were fed high protein primate chow (Purina Monkey Chow, St. Louis, MO) supplemented with fruit and multivitamins to maintain target body weights. Behavioral experiments were conducted at the same time each day, using protocols that were approved by the Institutional Animal Care and Use Committee at McLean Hospital. All subjects were housed in the McLean Hospital Animal Care Facility (licensed by the US Department of Agriculture) in accordance with guidelines provided by the Committee on Care and Use of Laboratory Animals of the Institute of Laboratory Animals Resources, Commission on Life Sciences, National Research Council [27]. All subjects were drug naïve prior to experimentation and were implanted with a chronically indwelling intravenous (i.v.) catheter [28] that was protected by a custom-fitted nylon vest (Lomir Biomedical Inc., USA).

### Apparatus

During experimental sessions, subjects sat in customized Plexiglas chairs that were enclosed in ventilated, sound- and light-attenuating chambers; white noise was present at all times during experimental sessions to mask extraneous sounds. Each subject faced a front panel consisting of red stimulus lights, two response levers, and a receptacle (1/4” deep and 1” diameter) into which milk could be delivered. Presses on either lever with a force of >0.2 N were recorded as a response. A 10-RPM motor-driven syringe pump (Med-Associates, Inc., Georgia, VT, Model PHM-100-10) outside the chamber could be operated to deliver milk via Tygon tubing to the receptacle (0.15 ml in 0.84 s; 60-ml syringe) or i.v. injections of vehicle or drug solution via a venous catheter (0.1 ml in 0.84 s; 30-ml syringe). Experimental variables and data collection were controlled and recorded via interfacing equipment and operating software (MED-PC, MedState Notation; Med Associates Inc., St. Albans, VT).

### Training

All subjects were trained on concurrent second-order fixed-ratio (FR) 3, FR5 schedules [FR3(FR5:S)] of reinforcement. Levers were counterbalanced between subjects as either an injection lever or milk lever, and the conditions associated with each lever remained constant throughout the study. Under these conditions, the availability of reinforcement in daily 90-min sessions was signaled by the illumination of a red stimulus light, and the completion of an FR5 produced a 0.2-s change from red to green stimulus lights (stimulus flash; S). Completion of 3 repetitions of an FR5:S produced a yellow stimulus flash accompanied by the delivery of 0.1 ml milk or water in the receptacle, or 0.1 ml injection of drug or saline through the i.v. catheter. Only consecutive responses culminating in the completion of the FR3(FR5:S) schedule were reinforced- i.e., a response on one lever reset the response requirement on the other lever. Each reinforcer delivery was followed by a 45-s time-out (TO) period in which all stimulus lights were turned off and responding had no programmed consequences. During training, subjects started under an FR1(FR1:S);TO1s of reinforcement, with only two possible conditions: 0.032 mg/kg/inj oxycodone vs water or saline vs 20% milk. Training was conducted under a double-alternation schedule- i.e., two consecutive days of 0.032 mg/kg/inj oxycodone vs water followed by two consecutive days of saline vs 20% milk, ad libitum. The schedule of reinforcement was systematically increased simultaneously on both levers (to prevent lever bias) until the terminal schedule of FR3(FR5:S);TO45s was reached. Once at the terminal schedule of reinforcement, subjects continued to train until they responded >80% on the drug lever under the 0.032 mg/kg/inj vs water condition, and >80% on the milk lever under the saline vs 20% milk condition for 4 out of 5 days. Once these training criteria were met, testing with various oxycodone doses and/or milk concentrations began.

### Self-Administration (Drug-Taking)

Self-administration test sessions were identical to those in the training sessions- i.e., 90-min sessions in which various doses of oxycodone and various concentrations of milk were concurrently available under FR3(FR5:S);TO45s schedules of reinforcement. There were no limitations on the maximum amount of drug and/or liquid deliveries that could be earned within the 90-min experimental sessions. Intravenous doses of oxycodone studied ranged from 0.001-0.1 mg/kg/inj, whereas milk concentrations included 0% (i.e., water), 5%, 10%, 20%, and 30%. Under each milk condition, doses of oxycodone were studied from where responding was predominantly distributed to the milk lever (<20% %ILR) up to a dose in which responses were predominantly distributed to the drug lever (>80% %ILR). Combinations of an oxycodone dose with a particular milk concentration were randomized within and between subjects. However, an individual subject was studied under the same oxycodone and milk conditions until the following stability criteria were met: 1) <15% change in %ILR for 2 out of 3 days; and 2) <15% change in total drug injections earned for 2 out of 3 days. The oxycodone dose-response function per milk concentration was determined to be complete if the prototypical inverted U-shaped curve was observed in the total number of injections earned per session- i.e., at least one dose with injections/session not significantly greater than saline followed by at least one subsequent dose greater than saline, and another large dose not different from saline. As a result of that crierion, 0.001 mg/kg/inj oxycodone was studied in females, but not males, when 10% milk was available. Likewise, 5% milk availability was only presented to males during oxycodone self-administration in an attempt to find levels of oxycodone self-administration similar to when water was available. Because self-administration of oxycodone was not appreciably different when either water or 10% milk were available, the 5% milk condition was not tested in females.

### Oxycodone-Primed Reinstatement (Drug-Seeking)

Reinstatement tests were 90-min in length, and although different concentrations of milk were randomly available between sessions, only saline injections were available during reinstatement tests. Reinstatment tests were not conducted with a multi-day period of extinction responding, but instead were conducted on days in which saline was administered (c.f. [13,29]). Both saline and milk were concurrently available under an FR3(FR5:S);TO45s schedule of reinforcement. Ten minutes prior to reinstatement tests, 0.32 mg/kg oxycodone was administered to the subject i.m. This dose of oxycodone was selected because previous studies have shown that this dose of oxycodone does not completely disrupt food-maintained operant behavior and can significantly reinstate opioid-seeking behavior [29]. Milk concentrations that were studied were 0%, 10%, 20%, and 30%, and reinstatement tests were conducted only once. No more than 2 reinstatement tests were conducted each week, and between tests, training was conducted with either 0.032 mg/kg/inj oxycodone vs water or saline vs 20% milk.

### Drugs and dosing procedures

Oxycodone hydrochloride was obtained from Sigma-Aldrich (St. Louis, MO) and dissolved in 0.9% NaCl solution and sterilized via filtration through a 0.22 μm filter (Millex-GV; Cork, Ireland). Doses are expressed in terms of the free base. Various concentrations of sweetened condensed milk (Casa Solano, Sysco, Plymptom, MA) were made fresh daily by diluting with water. The order of doses and milk concentrations studied in self-administration and reinstatement procedures were tested in a mixed order across subjects.

### Data Analysis

For both self-administration and reinstatement procedures, the number of reinforcer deliveries and responses emitted on each lever were recorded throughout the session. The percentage of injection-lever responding (%ILR) was calculated by dividing the number of responses emitted on the drug-lever by the total number of responses on both levers. Data from repeated determinations of self-administration studies (average 2-3 determinations/test) were averaged for individual subjects. Experimental results are presented as group means (±SEM).

For self-administration tests, total milk deliveries as a function of concentration when saline was available was compared using a repeated-measures analysis of variance (ANOVA) with Dunnett’s multiple comparison test. Dose-response functions for oxycodone generated with varying milk concentrations were analyzed using an F-ratio test. Straight lines were fitted to the linear portion of the dose-response functions and were utilized to examine changes in the potency of self-administered oxycodone as a function of available milk concentration. This was conducted using a linear regression when more than two points were available but were otherwise calculated by interpolation to estimate the unit dose that would result in the allocation of 50% responding to the injection-lever (ED_50_). In the case of injections earned per session, the ED_50_ was calculated by determining the dose that produced half of the maximal injections earned per session. Potency ratios were calculated by dividing the ED_50_ value of the least potent oxycodone curve by the most potent oxycodone curve. Because the %ILR did not drop below 50% when water was available, potency ratios were determined by comparing ED_75_ values. A repeated-measures two-way ANOVA with Sidak’s post-hoc test examined whether the number of injections per session varied as a function of dose or sex, or if the number of milk deliveries varied as a function of milk concentration or sex. The oxycodone dose that engendered the greatest amount of inj/session under each milk condition was determined per individual subject- i.e., the dose of oxycodone with the most responding may not have been the same in all subjects. The peak inj/session/milk concentration were averaged per sex, and a repeated-measures one-way ANOVA with a Dunnett’s Multiple Comparison’s test examined whether the peak number of oxycodone injections earned per sex varied as a function of concurrently available milk concentrations. A two-tailed t-test with a Bonferroni’s correction for multiple comparisons compared the peak number of oxycodone injections for each sex with different milk availabilities.

For reinstatement tests, total %ILR, cumulative drug lever responses, and cumulative milk lever responses were plotted in 30-min bins, and compared using two-way ANOVA, with time and milk concentration factors for analysis, followed by Tukey’s multiple comparison post-hoc test. Cumulative drug lever responses were normalized to the average baseline responses at each time point and the areas under the curve were calculated for the normalized time response function. A repeated-measures two-way ANOVA with a Sidak’s post-hoc test compared whether total number of saline and/or liquid deliveries varied when different concentrations of milk were available. Significance was set at p<0.05.

## Results

### Responding for Oxycodone or Milk Alone

Self-administration of a range of oxycodone doses was first studied with water as the available alternative (Figure 1, left panel) and responding for several milk concentrations was studied with saline as the response-contingent injection (Figure 1, right panel). When water was the available alternative, oxycodone generated the characteristic inverted-U shaped curve observed in i.v. self-administration studies under second-order schedules of reinforcement [29]. The number of injections earned per session varied as a function of dose (F_3,18_=5.1, p=0.01), but not sex. At least one dose of oxycodone was found to be statistically greater than saline in both males and females when each milk concentration was available. When saline was the response contingent injection, increasing the concentration of milk from water to 30% increased the number of milk deliveries per session from 9.5 to 87 in males and 25 to 85 in females with no significant differences between males and females regarding total number of milk deliveries. There was a significant effect of milk concentration on milk deliveries/session when saline was available in both males (F_4,11_=7.5, p=0.0037) and females (F_4,10_=11, p=0.0012), with post-hoc analysis indicating that 10, 20, and 30% milk were significantly greater than water.

**Figure 1.**
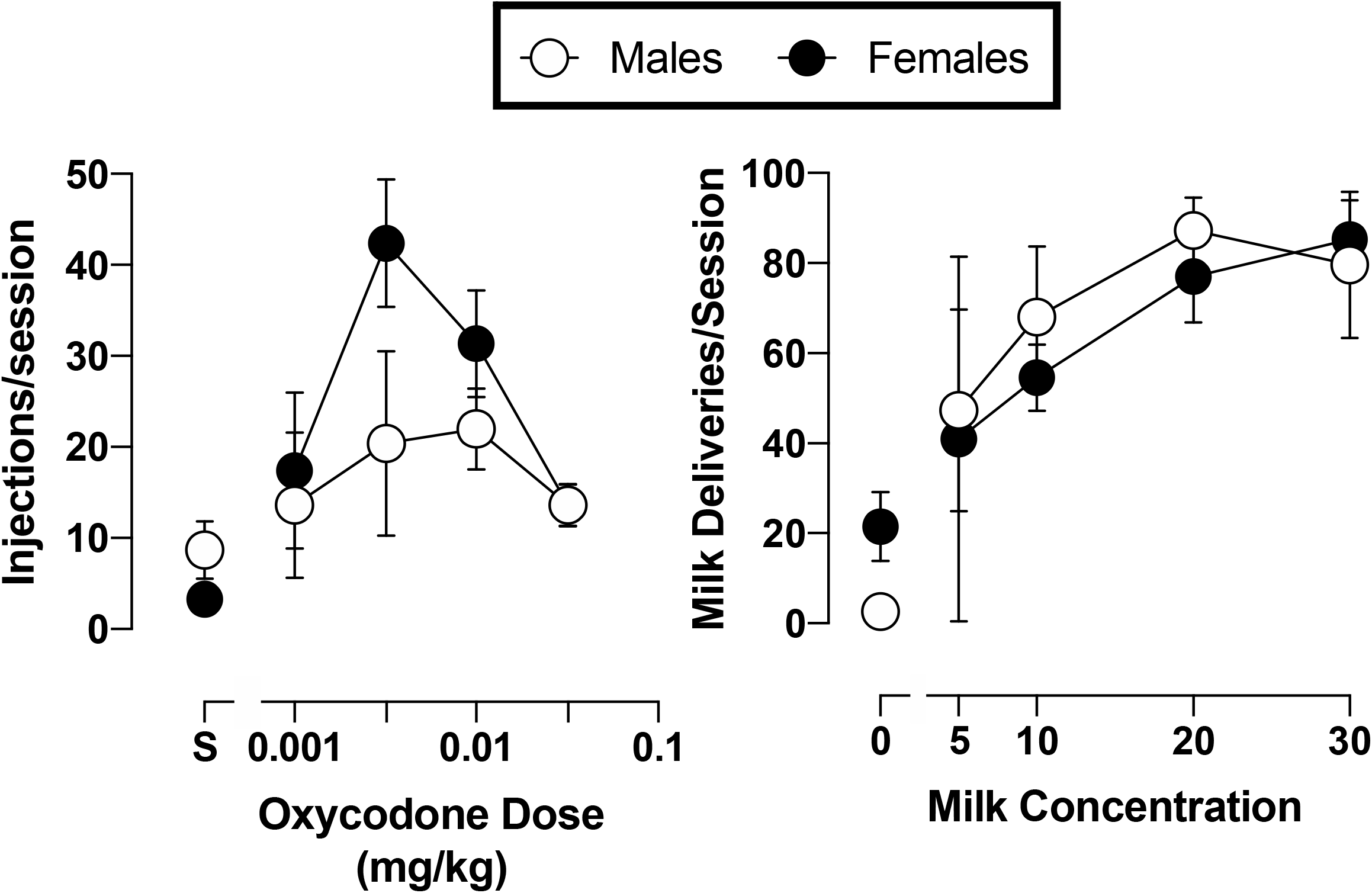
Allocation of behavior on the drug-paired lever when water is available (left) and the number of milk deliveries earned when saline is available (right). Males: open circles. Females: closed circles. (left) ordinate: %ILR; abscissa: oxycodone dose in mg/kg. (right) ordinate: number of milk deliveries earned per session; abscissa: milk concentration.

### The Effect of Alternative Reinforcer Magnitude on Oxycodone Self-Administration in Males

When oxycodone was available for self-administration, increasing the concentration of the alternative reinforcer from water to 30% milk significantly decreased the potency of oxycodone to function as a reinforcer in males (Figure 2, top panels). Regardless of concentration, the number of saline injections earned per session averaged fewer than two when milk was available. Milk availability significantly shifted the ascending limb of the oxycodone injections/session curve rightward and downward (F_8,29_=2.3, p=0.048; Figure 2, top left panel). The availibility of 10, 20, and 30% milk shifted the injections/session dose-response function 16-, 62-, and 22-fold rightward, respectively (Table 1). All changes in potency of oxycodone injections/session were selective for the ascending limb of the oxycodone dose-response function, with no effects on the descending limb.

**Figure 2.**
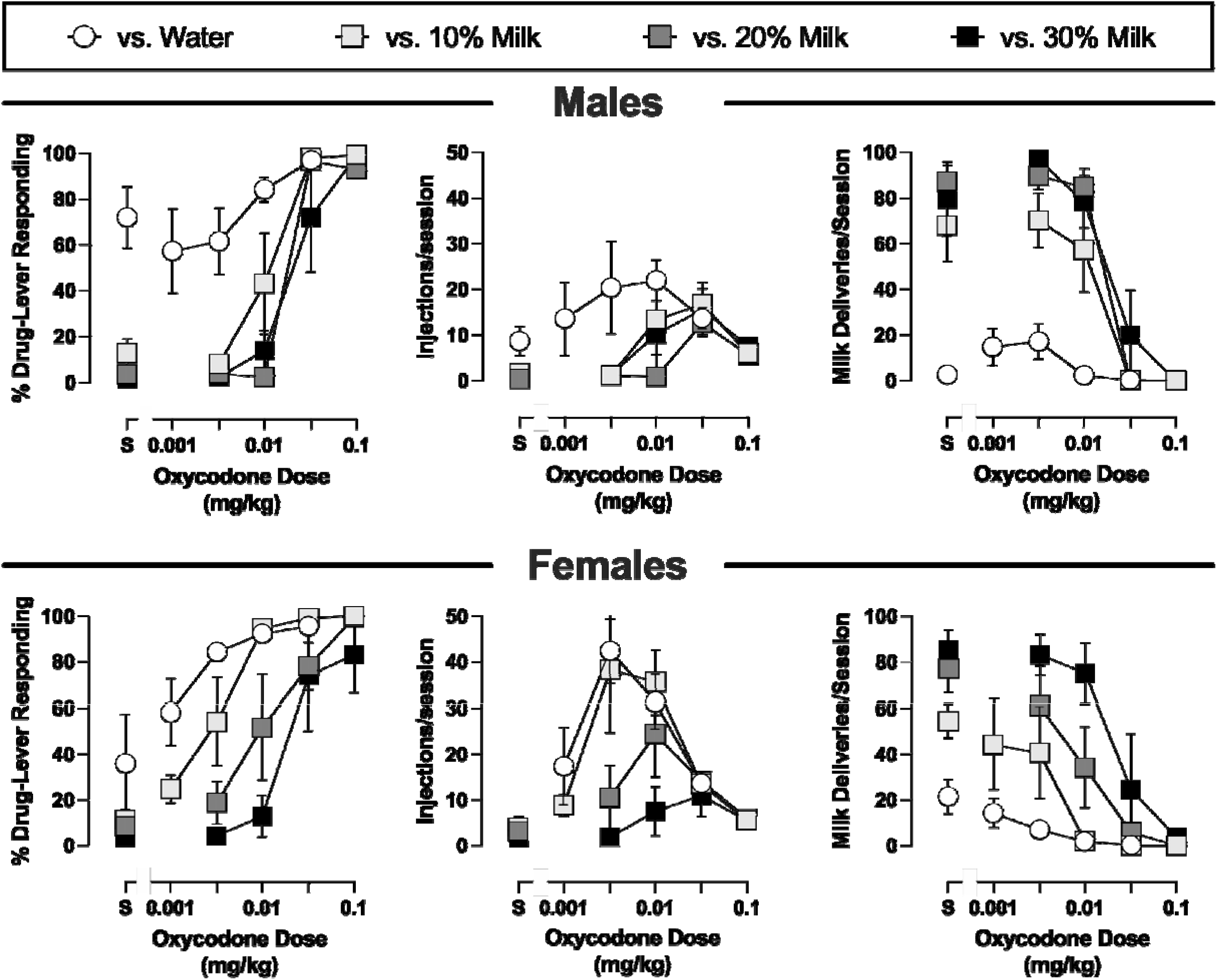
Self-administration of oxycodone in males (top panels) and females (bottom panels) when different concentrations of milk are available. Left panels: ordinate- drug injections earned per session; middle panels: ordinate- milk deliveries earned per session; right panels: ordinate- %ILR. All panels: abscissa- oxycodone dose in mg/kg.

**Table 1.** Analysis of self-administration and reinstatement of oxycodone in males and females as a function of the availability of alternative.

| Self-Administration | %ILR | Drug Injections | Milk Deliveries | %ILR ED<sub>75</sub><br>(mg/kg) | %ILR PR<br>(95% CI) | Inj ED<sub>50</sub><br>(mg/kg) | Inj PR<br>(95% CI) |
| --- | --- | --- | --- | --- | --- | --- | --- |
| *Males* |  |  |  |  |  |  |  |
| Water |  |  |  | 0.008 (0.0037 - 0.017) |  | 0.0012 (0.00032 - 0.051) |  |
| 5% | F<sub>2,11</sub>=7.4, p=0.009 | F<sub>2,11</sub>=2.7, 9=0.11 | F<sub>2,9</sub>=1.5, p=0.28 | 0.018 (0.0028 - 0.12) | 2.2 (0.8-6.6) | 0.008 (0.004 - 0.018) | 6.8 (0.1 - 358) |
| 10% | F<sub>2,14</sub>=4.4, p=0.032 | F<sub>2,14</sub>=1.2, p=0.33 | F<sub>2,13</sub>=5.0, p=0.025 | 0.035 (0.0036 - 0.34) | 4.4 (1.3-15) | 0.011 (0.0044 - 0.027) | 16 (1.1 - 211) |
| 20% | F<sub>2,14</sub>=32, p<0.0001 | F<sub>2,14</sub>=2.8, p=0.093 | F<sub>2,14</sub>=21, p<0.0001 | 0.048 (0.041 - 0.056) | 6.5 (3.4-12) | 0.027 (0.018 - 0.041) | 62 (1.1 - 342) |
| 30% | F<sub>2,15</sub>=5.7, p=0.015 | F<sub>2,14</sub>=1.9, p=0.19 | F<sub>2,15</sub>=18, p=0.0001 | 0.062 (0.0088 - 0.44) | 7.9 (2.2-28) | 0.014 (0.0043 - 0.043) | 22 (1 - 521) |
| *Females* |  |  |  |  |  |  |  |
| Water |  |  |  | 0.0025 (0.0022 - 0.0028) |  | 0.0012 (0.00055 - 0.0026) |  |
| 10% | F<sub>2,14</sub>=2.4, p=0.13 | F<sub>2,14</sub>=0.85, p=0.45 | F<sub>2,9</sub>=2.3, p=0.15 | 0.0029 (0.0024 - 0.0036) | 1.5 (1.2-1.9) | 0.0015 (0.00061 - 0.0036) | 1.3 (0.6 - 3.1) |
| 20% | F<sub>2,15</sub>=9.8, p=0.0019 | F<sub>2,12</sub>=3.4, p=0.066 | F<sub>2,13</sub>=5.7, p=0.017 | 0.011 (0.006 - 0.022) | 11 (5.1-25) | 0.0038 (0.0009 - 0.016) | 6.6 (2 - 21) |
| 30% | F<sub>2,13</sub>=4.8, p=0.028 | F<sub>2,9</sub>=6.5, p=0.018 | F<sub>2,13</sub>=7.3, p=0.0077 | 0.028 (0.0056 - 0.14) | 29 (4.8-171) | 0.0093 (0.0022 - 0.04) | N/A |
| Reinstatement | Time | Milk Concentration | Interaction | Water AUC<br>(± SEM) | 10% AUC<br>(± SEM) | 20% AUC<br>(± SEM) | 30% AUC<br>(± SEM) |
| *Males* |  |  |  |  |  |  |  |
| %ILR | F<sub>2,9</sub>=9.6, p=0.0059 | F<sub>3,24</sub>=12, p<0.0001 | F<sub>6,24</sub>=1.7, p=0.16 | 9119 ± 1489 | 9567 ± 1447 | 8848 ± 3073 | 4831 ± 1416^ |
| Drug Injections | F<sub>2,9</sub>=3.9, p=0.06 | F<sub>3,24</sub>=4.5, p=0.01 | F<sub>6,24</sub>=1.2, p=0.35 |  |  |  |  |
| Milk Deliveries | F<sub>2,9</sub>=17, p=0.0009 | F<sub>3,24</sub>=15, p<0.0001 | F<sub>6,24</sub>=4.6, p=0.0029 |  |  |  |  |
| *Females* |  |  |  |  |  |  |  |
| %ILR | F<sub>2,9</sub>=7, p=0.015 | F<sub>3,27</sub>=5.1, p=0.0063 | F<sub>6,27</sub>=1.3, p=0.31 | 11325 ± 4531 | 7185 ± 1489^ | 6490 ± 2612^ | 6602 ± 1543^ |
| Drug Injections | F<sub>2,9</sub>=1.7, p=0.23 | F<sub>3,27</sub>=3.6, p=0.026 | F<sub>6,27</sub>=1.2, p=0.91 |  |  |  |  |
| Milk Deliveries | F<sub>2,9</sub>=7.4, p=0.013 | F<sub>3,27</sub>=10, p=0.0001 | F<sub>6,27</sub>=3.7, p=0.0087 |  |  |  |  |

reinforcers of varying magnitude. ED<sub>75</sub>- effective dose 75%; ED<sub>50</sub>- effective dose 50%; PR- potency ratio; AUC- aread under the curve; SEM- standard error of the mean
^Significantly different from Water AUC (p<0.05)

The total number of milk deliveries earned when various doses of oxycodone were available also significantly varied as a function of milk concentration (F_8,33_=6.0, p<0.0001; Figure 2, top middle panel). When saline was available, milk increased the total liquid deliveries in a concentration dependent manner with maximal deliveries reaching, on average, 97 per session at the highest milk concentration (i.e., 30%). Increasing the dose of oxycodone produced significant decreases in milk intake which is reflected as an increase in the distribution of behavior onto the oxycodone lever, i.e., %ILR (Figure 2, top right panel; Table 1). Milk concentrations of 10, 20, and 30% each produced up to an 8-fold decrease in the potency of oxycodone to function as a reinforcer in males (see Table 1).

### The Effect of Alternative Reinforcer Magnitude on Oxycodone Self-Administration in Females

In female subjects, saline injections/session never averaged more than 3 when milk was available. Similar to males, increasing the concentration of milk available under choice conditions significantly shifted the ascending limb of the oxycodone dose-response function rightward and downward (F_6,26_=4.3, p=0.0036; Figure 2, bottom left panel). Although there was no significant difference in the ascending limb of the oxycodone curve when water or 10% milk was available, increasing the available milk concentration to 20% shifted the ascending limb 6.6 (95% CI:2.0-21)-fold to the right. Because the slope of the ascending limb of the oxycodone dose-response function was significantly different when 30% milk was available compared to when water or 10% milk were available, a potency ratio could not be calculated. However, 30% milk shifted the oxycodone dose-effect function downward and rightward compared to water (F_2,9_=6.5, p=0.018) and 10% milk (F_2,14_=7.4, p=0.0064) availability. The shift in the ascending limb of the oxycodone dose-response function with various milk concentrations paralleled an increase in milk deliveries earned per session (F_6,25_=4.0, p=0.005; Figure 2, bottom middle panel). There was no significant difference in milk deliveries per session between water and 10%, but there was a significant difference between water and 20% or 30% milk (Table 1). Furthermore, increasing the milk concentration nearly doubled the number of milk deliveries earned per session (Figure 2b).

As in male subjects, increasing the milk concentration (20% and 30%) significantly decreased the potency of oxycodone to increase %ILR (Table 1; Figure 2, bottom right panel). This suggests that the differences observed in %ILR were heavily influenced by the rightward and upward shifts in the milk deliveries/session observed when the milk concentration was increased from water (Figure 2, bottom panels). In contrast to males however, females appeared to be more sensitive to the reinforcing effects of oxycodone as a unit dose of 0.0032 mg/kg/inj oxycodone produced appreciably greater %ILR and drug/inj/session when water or 10% milk were available compared to saline (Figure 2, bottom panels).

### The Effect of Milk on Oxycodone-Primed Reinstatement in Males

Figure 2 shows that saline pretreatment did not engender significant oxycodone-lever responding in males when either of the milk concentrations were available (see points above “S” on x-axis). Therefore, a saline pretreatment did not elicit resinstatement behavior in males. Figure 3 (top panels) shows that oxycodone (0.32 mg/kg) elicited responses primarily on the drug-associated lever (>90%) throughout sessions when water was available. In constrast, 30% milk availibility shifted the distribution of responding from the drug-associated lever (~75%) during the first 30-min of the session to the milk-associated lever during the final 60-min (Figure 3, top right panel). While %ILR at the 90-min timepoint was decreased to <40% when 10 and 20% milk were available, this effect was largely driven by increased responding for milk and not a reduction in drug lever responses (Figure 3; top right panel; see Table 1).

**Figure 3.**
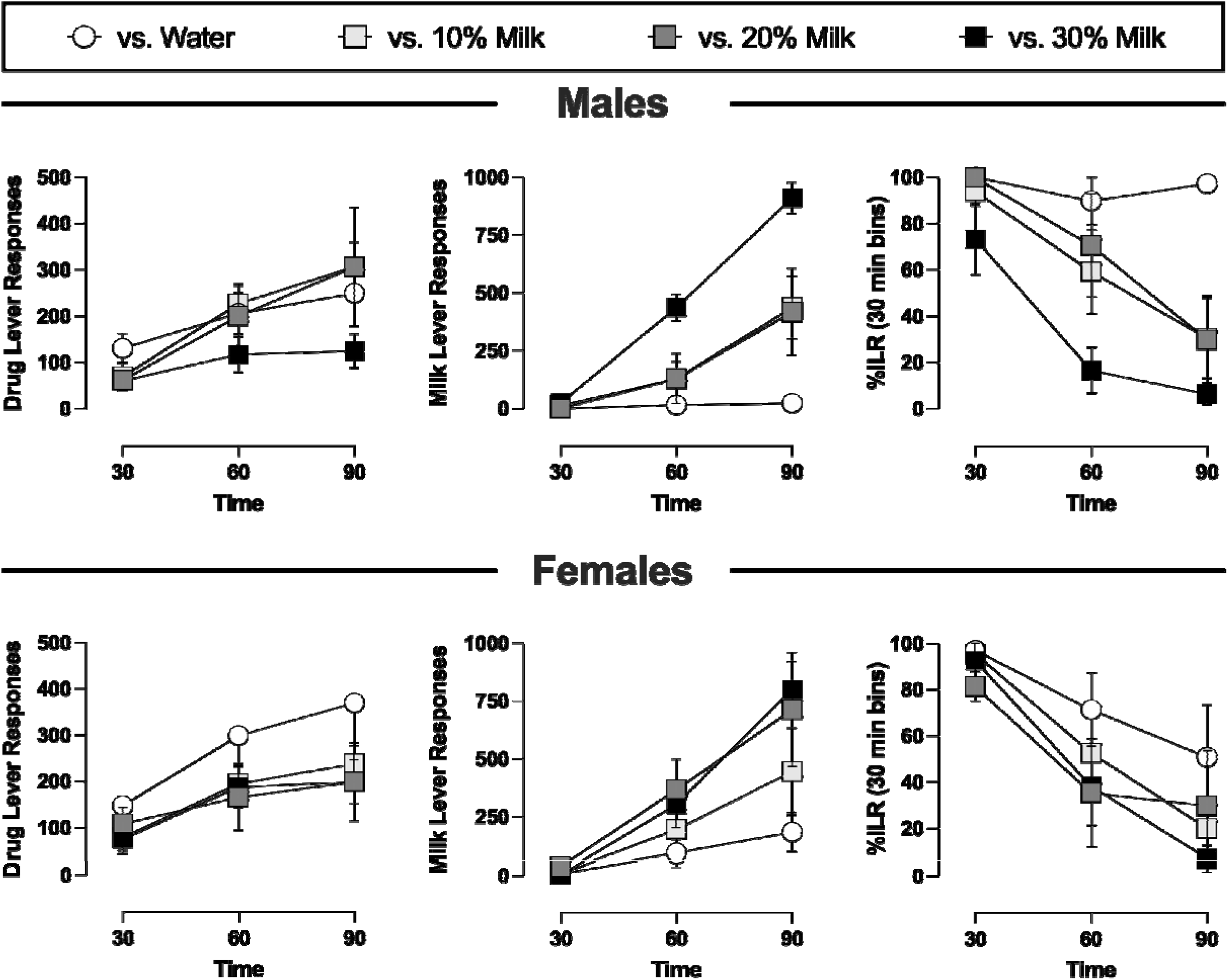
Oxycodone drug-seeking in males (top panels) and females (bottom panels) when different concentrations of milk are available. All panels: abscissa- time in 30-min bins. Left panels: ordinate- cumulative drug-paired lever responses; middle panels: ordinate- cumulative milk-paired lever responses; right panels: ordinate- %ILR in each 30-min bin.

### The Effect of Milk on Oxycodone-Primed Reinstatement in Females

Similar to what was observed in males, pretreatment with saline did not engender responding on the oxycodone-associated lever (see Figure 2), indicating that saline does not produce reinstatement in females. Unlike in males however, the distribution of behavior under water vs saline conditions following a 0.32 mg/kg oxycodone pretreatment shifted from responding predominantly on the drug-paired lever to the milk-paired lever over time, as evidenced by an approximately 50% decrease in %ILR between the first and last 30-min bins (Figure 3, bottom right panel). The availability of milk as an alternative produced a greater and more rapid shift in allocation of behavior (Table 1) with each of the milk concentrations producing a significant decrease in cumulative drug lever responses relative to water (Figure 3, bottom left panel).

### Comparison of Self-administration and Reinstatement in Males and Females

To better understand the relationship between sex and the availability of an alternative reinforcer on oxycodone’s abuse-related behavioral effects, the potency of oxycodone to function as a reinforcer (Figures 4, left; Table 1 top) or induce reinstatment (Figures 4, right; Table 1 bottom) during the availibility of each milk concentration was compared between the sexes. These analyses found that, when compared to water, the potency ratio of oxycodone at the different milk concentrations had differing profiles in males and females for self-administration and reinstatement. In self-administration tests, the various concentrations of milk studied produced a relatively invariant attenuation of oxycodone’s potency as a reinforcer (i.e., increase %ILR) in males, which was reflected as modest rightward shifts relative to water (4.4 [1.3-15]-, 6.5 [3.4-12]-, and 7.9 [2.2-28]-fold at 10, 20, or 30% milk, respectively). In contrast, the availibility of milk produced a concentration-dependent shift in oxycodone potency in female subjects, with the potency decreasing from 1.5 (1.2-1.9)-, 11 (5.1-25)-, and 29 (4.8-171)-fold at 10, 20, and 30% milk availability, respectively, compared to water. The potency of oxycodone was similar between the sexes when water and 30% milk was available but was 5.1 (2.4-11)- and 2.1 (1.1-4.0)-fold less potent in males relative to females when 10% and 20% milk was available, respectively. As such, all concentrations of milk effectively changed the distribution of behavior in males, whereas the lowest milk concentration was ineffective in females; the higher milk concentrations produced a greater rightward shift in %ILR in females. The peak number of oxycodone injections earned per session did not significantly differ between males and females except when 10% milk was available (T_6_=2.34, p=0.029), where males received on average 19 inj/session and females received on average 43 inj/session. Peak number of oxycodone injections earned per session was significantly lower in males when 10%, 20%, and 30% milk were available compared to water, whereas peak injections were only significantly lower in females when 30% milk was available (Figure 4, left).

**Figure 4.**
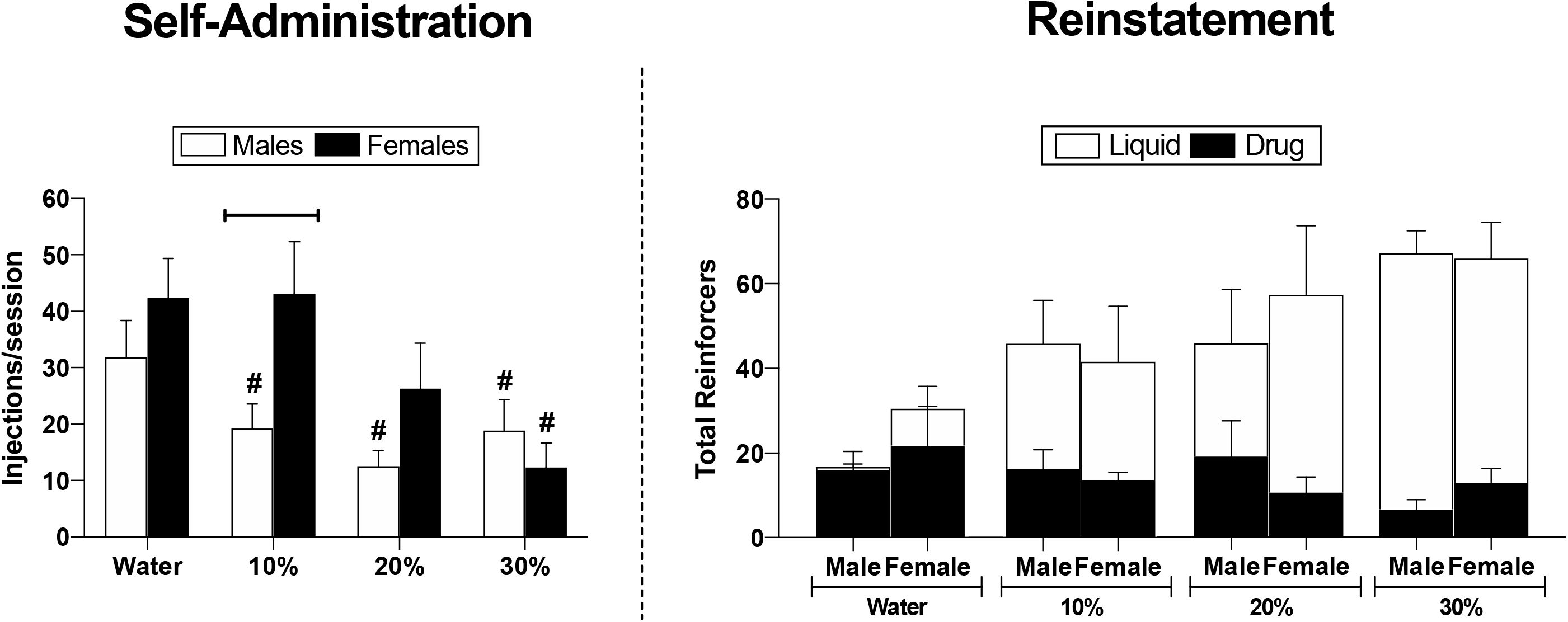
Comparison of the peak number of oxycodone injections/session relative to available milk concentration in the drug-taking studies (left panels), and the effects of the various milk concentrations in the drug-seeking studies (right panels). (Left) The number of oxycodone injections/session when 10, 20, and 30% milk was available compared to water. Ordinate: injections/session; abscissa: milk concentration. (Right) A comparison of the total number of oxycodone and liquid reinforcers earned per session between females and males when water, 10, 20, and 30% milk was available. Ordinate: total reinforcers earned per session; abscissa: milk concentration.

In reinstatement tests, 30% milk significantly decreased drug-lever responding in both males and females, but 10 and 20% milk were only effective in females (Figure 4, right; Table 1). The AUC when water and 30% milk were available was greater in females, but was greater in males when 10 and 20% milk were available (Figure 4, right). However, although the total number of lever presses, and as a result, AUC, differed between males and females, the total number of reinforcer deliveries did not signficantly differ between the sexes. This counterintuitive finding most likely reflects a tendency of females to distribute reponses on both levers without completing the terminal schedule with a greater frequency than males. Despite that, saline deliveries decreased, and liquid deliveries increased in all subjects as a function of increasing milk concentrations.

## Discussion

The use of non-drug alternative reinforcers has long been utilized as a component of therapeutic interventions for the management of substance use disorder [4,5,6,30]. For example, under controlled laboratory conditions in humans, when presented with a choice between various amounts of money or drug dosages, intake of cocaine, nicotine, alcohol, and heroin significantly decreases compared to when no alternative reinforcer is present [7,8, 9,10]. As a result, such interventions (e.g., money, job) have been incorporated into clinical treatment programs such as contingency management and voucher programs [4,5,11]. Preclinical studies in both nonhuman primates and rodents demonstrate that the availability of a palatable alternative reinforcer (e.g., palatable foods, exercise, enriched environments, etc.) decreases intake of various drugs of abuse [12,13,15,16]. The present results expand on those previous findings and demonstrate that the magnitude of the alternative reinforcer can systematically decrease the reinforcing potency of oxycodone and decrease relapse-related drug-seeking behavior in both male and female nonhuman primates.

The magnitude of a reinforcer in schedule-controlled behavioral assays can influence the rate of operant responding and the number of reinforcers earned, which likely reflects relative reinforcing strength of various reinforcers. For example, using a progressive ratio schedule of reinforcement, previous studies have demonstrated that breakpoint for 3 sucrose pellets is significantly greater than for one sucrose pellet [32]. Likewise, breakpoints for abused drugs, like oxycodone, systematically increase as a function of increased unit dose [33]. These principles of reinforcement were readily observed in the current study; when saline was concurrently available, the number of sweetened condensed milk reinforcers earned per session increased as a function of milk concentration, consistent with the hypothesis that larger concentrations of milk are more reinforcing than lower. However, oxycodone dose-dependently decreased the number of milk reinforcers earned regardless of concentration, which was reflected by the %ILR. This finding is consistent with previous studies describing the effects of alternative reinforcers on self-administration of the monoaminergic ligands cocaine and procaine [16]. Here, we extend this work to show that the potency of the μ-opioid agonist oxycodone to preferentially engender drug-lever responding was systematically attenuated by larger magnitudes of a concurrently available food reinforcer. Furthermore, the peak number of oxycodone injections earned per session progressively decreased relative to the concentration of the concurrently available milk, suggesting that under these conditions, the presence of a competing non-drug alternative decreases the efficacy of oxycodone to function as a reinforcer.

Although the impact of alternative reinforcer magnitude on drug self-administration has been previously investigated, there is a paucity of information regarding the effects of alternative reinforcers on drug-primed reinstatement (i.e, drug-seeking). Contingency management studies have shown that alternative reinforcers can decrease relapse rates in humans [34], and a recent study in rodents found that asynchronous access to wheel-running exercise decreases heroin-seeking behaviors in rats [35]. However, the extent to which reinforcer magnitude differentially impacts the efficacy of treatment programs that utilize alternative reinforcers is unclear. The current report is the first preclinical study to systematically investigate the impact of concurrently available nondrug alternative reinforcers on opioid-seeking behaviors. The data presented here suggest that the availability of a palatable food alternative reinforcer attenuates oxycodone-primed reinstatement in squirrel monkeys. These findings are consistent with the self-administration findings inasmuch as milk availability appears to decrease the efficacy of oxycodone. Importantly, in the absence of an alternative reinforcer- i.e., when water is available-behavior is primarily directed toward drug-seeking, a finding that translates to the clinic insofar as relapse rates increase when patients are no longer, or were never, involved in a treatment program that facilitated access to alternative reinforcers [4,36].

There is a growing appreciation for the role that sex as a biological variable may play in the abuse-related effects of drugs from various pharmacological classes, including prescription opioids [37]. Previous clinical and preclinical findings suggest that oxycodone, and other opioids, have greater efficacy in males compared to females. A compelling argument for this hypothesis is that both spontaneous and precipitated withdrawal symptoms in humans, nonhuman primates, and rodents are greater in magnitude in males [38]. The present findings are consistent with human and rodent studies inasmuch as females self-administered oxycodone at lower doses, and responded for a greater number of injections at the peak dose, compared to males [25] when 10% milk was available. Typically, higher doses of a drug reinforcer produce fewer total injections/earned per session, reflecting that less behavior is needed to maintain steady states of drug reinforcement. Consequently, more schedule completions are required for lower drug doses, which can be seen as having lesser functional efficacies, to maintain a similar steady state. Following that logic, greater peak numbers of injections/session observed in females may be a reflection of a less efficacious drug. However, it is important to recognize that a difference in the peak number of injections/session earned between males and females was only observed when water or 10% milk was the available alternative. This may suggest that although the efficacy of oxycodone is less in females, its potency is greater. As a result, females are more likely to detect the presence of oxycodone even when an alternative reinforcer is present, and furthermore, will more likely self-administer lower doses of oxycodone that are less likely to constrain the total number of injections/session. That lower doses of oxycodone were self-administered in females, but not males, under some conditions may suggest that females are more sensitive to the interoceptive effects of oxycodone. In support of this view, previous studies have demonstrated that μ-opioid agonists such as morphine and buprenorphine substitute for a morphine discriminative stimulus at lower doses in females compared to males [39]. Overall, this interpretation of the present results suggests that oxycodone is more potent, and less efficacious, in females compared to males. Although speculative, this hypothesis also may explain why both the adverse (e.g., nausea) and reinforcing effects (e.g., liking) have been described in the clinical literature as greater in females.

Our data show that when water was available during reinstatement studies, males responded on the drug-paired lever throughout the entire session, whereas the distribution of behavior in females shifted to the milk-paired lever within the session. This change in the distribution of responding in females may reflect a greater proficiency in detecting the presence, or absence, of oxycodone. However, despite that observation, the absolute number of saline injections/session, a metric of drug-seeking (relapse-related) behavior, did not quantifiably differ as a function of sex. Instead, the results from this study are consistent with the clinical literature in which alternative reinforcers are equieffective in decreasing the probability of relapse in males and females. Importantly, in the absence of alternative reinforcer- i.e., when water was available-the distribution of behavior in males and females was greatest on the drug lever. That drug-seeking was greater in the absence of alternative reinforcers is consistent with clinical reports of increased risk of relapse following the completion of behavioral treatment programs such as contingency management or voucher programs [4,36].

The biological basis for sex differences in behavioral responses to opioids has not yet been fully elucidated, however, the influence of gonadal hormones – both ovarian hormones and androgens – on endogenous opioid systems and opioid-induced behaviors is well documented (reviewed by [40,41,42,43,44]). In addition to direct regulation of opioid receptors and/or endogenous peptides, another possible mechanism for sex differences is that the pharmacokinetics of oxycodone differs between males and females. Previous studies have shown that although oxycodone plasma levels are similar between males and females following oxycodone administration [23,45,46], oxycodone levels in the brain are greater in males compared to females [23,47]. Further, a recent study found that, following cytochrome P450 inhibition, female rats in diestrus had smaller increases in brain oxycodone concentrations and a smaller decrease in brain oxymorphone/oxocodone ratios compared to male rats or female rats in estrus [46] suggesting an interaction between gonadal hormone levels and pharmacokinetics. Additional studies are needed to provide insight into potential hormonal mechanisms involved in the effects described here.

The results from this study may have translational relevance. As OUD treatment programs such as contingency management and voucher programs utilize alternative reinforcers as a means for reducing drug-taking and preventing relapse, the results of the current study suggest that a larger incentive may be needed to curtail ongoing drug use in females. However, there are important limitations that should be considered. First, the present study used oxycodone and the generality of these results to other opioid drugs with different pharmacokinetic profiles or mu-receptor efficacy (i.e., heroin, fentanyl, buprenorphine) is unknown. Next, as the hormonal status of females used in this study was not monitored, the role of menstrual cycle phase and whether it may alter choice behavior or sensitivity to alternative reinforcer availability remains to be determined. Finally, access to treatment programs such as contingency management remains a significant barrier for many patients from underrepresented population and, increasing access to such programs is an important step in translating preclinical findings to clinical practice. Despite these limitations, the findings also highlight the utility of well-controlled choice self-administration procedures in evaluating behavioral and pharmacological factors which may have utility in the management of opioid use disorders. Overall, results from this study suggest that the magnitude of an alternative reinforcer can systematically alter both drug-taking and -seeking behavior of oxycodone in male and female nonhuman primates.

## Data Availibility Statement

Data will be made available to qualified individuals upon request to the corresponding author

## Acknowledgements

The authors would like to thank Erin Conley and Bryan Carlson for their technical assistance, and Dr. Roger Spealman for his comments on an earlier version of this manuscript.

## Author Contributions

Experimental Design: FBM and SJK

Conducted Experiments: FBM

Data Analysis: FBM and SJK

Writing of Manuscript: FBM, RGB, and SJK

